# The dilated cardiomyopathy E525K mutation stabilizes the cardiac myosin interacting-heads motif while activating the isolated motor domain

**DOI:** 10.64898/2026.10.02.756299

**Authors:** Ruchi Gautam, Arun Kumar Somavarapu, Kalen Z. Robeson, Matthew C. Childers, Michael Regnier, Jinghua Ge, Christopher M. Yengo, Roger Craig, Raúl Padrón

## Abstract

Mutations in β-cardiac myosin are a common cause of inherited cardiomyopathies. The dilated cardiomyopathy E525K mutation alters both thick filament regulation and motor activity, yet the structural basis for these effects has remained unclear. Here, we combined cryo-EM and MD simulations to determine how E525K affects the conformational landscape of cardiac myosin in both its autoinhibited interacting-heads motif (IHM) and isolated myosin head states. E525K-mutated myosin exclusively adopted a single S2 conformation in IHM, in contrast to the conformational heterogeneity observed in wild-type myosin. Structural and electrostatic analyses revealed that the mutation increased positive charge density at the blocked-head (BH) – S2 (subfragment-2) interface, strengthening interactions with negatively charged residues within S2. 3D variability analysis and MD simulations of the E525K IHM demonstrated reduced S2 mobility, consistent with strengthened BH – S2 interactions, and decreased conformational flexibility, providing a structural mechanism for stabilization of the IHM state. To investigate the isolated myosin head, we determined cryo-EM structures of wild-type and E525K subfragment-1 (S1). The mutation induced local conformational rearrangements in the activation-loop and Loop 3 region, the SH3-like domain, and the essential light chain of S1, revealing structural changes that support the enhanced motor activity previously reported for isolated E525K myosin motors. These findings show that E525K stabilizes the autoinhibited IHM through electrostatic stabilization of the BH – S2 interface while inducing structural changes consistent with activation of the isolated motor domain, providing structural insights into how a single DCM mutation differentially regulates cardiac myosin S1 and IHM structure and function.

## Introduction

Cardiac muscle contraction is tightly regulated by troponin-tropomyosin on the thin filament and the equilibrium between active (open) and autoinhibited (closed) states of myosin on the thick filament, ensuring that force generation is matched to physiological demand (1–3) (Fig. 1A). Structurally, myosin is composed of two heavy chains and two pairs of light chains (essential and regulatory, ELC and RLC) (Fig. 1C). The heavy chains dimerize through their C-terminal halves to form a coiled-coil tail, while their N-terminal regions fold into two globular heads, each containing a motor domain and a light chain binding domain (4)(Fig. 1C). In thick filaments, myosin molecules assemble into bipolar polymers in which the tails form the filament backbone, and the heads lie on the surface, where they can interact with actin filaments and generate force (5–7) (Fig. 1B).

**Fig 1.**
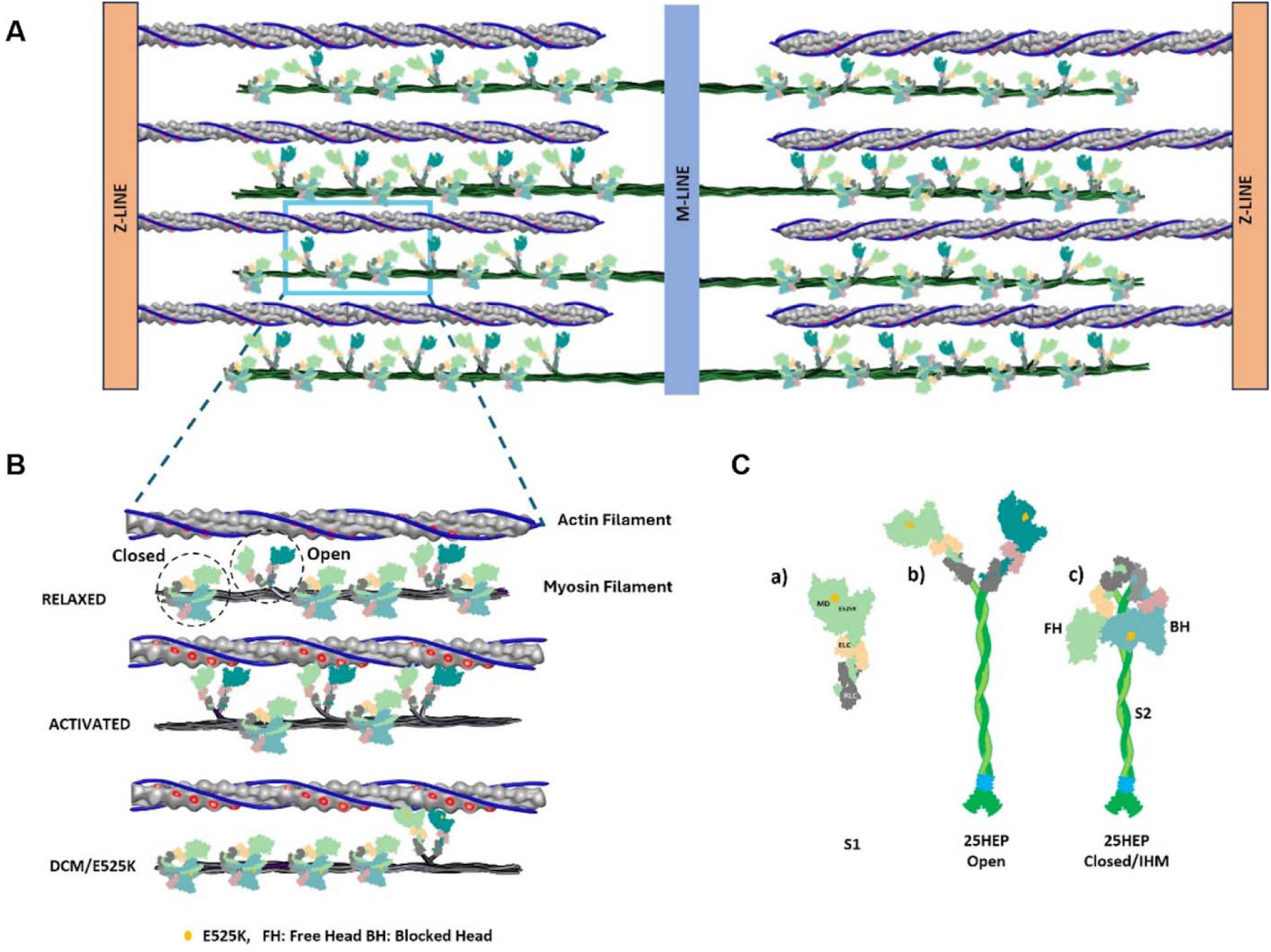
Schematic representation of the cardiac sarcomere and myosin constructs. **A.** The cardiac sarcomere showing thick filaments composed of myosin (green) overlapping with actin-containing thin filaments (grey) anchored at the Z-disc. Thick filaments are connected through their bare zones at the M-line. **B.** In relaxed muscle, myosin heads populate either the auto-inhibited closed/IHM (SRX) state or the open, disordered (DRX) state. Upon activation, myosin heads are released from the SRX population and engage actin filaments to generate contractile force. The DCM-associated E525K mutation stabilizes the folded IHM/SRX state, thereby reducing the number of myosin heads available for contraction, contributing to cardiac hypocontractility. **C.** Schematic representation of human β-cardiac myosin constructs used in this study, including isolated, single-headed S1 (a) and two-headed HMM (containing 25 heptads of S2 tail), in the open (b) and closed/IHM states (c). Constructs contain ELC and RLC light chains and a C-terminal GFP tag, while 25-hep additionally contain a GCN4 leucine zipper to promote two-headed dimerization.

Central to the cardiac contraction/relaxation cycle is the interacting-heads motif (IHM) of myosin, a closed off-state, relaxed conformation in which the heads fold back against the proximal S2 tail and interact with each other, thereby limiting their activity (5, 6, 8–13) (Fig. 1Cc). In this arrangement, the actin binding loop regions of the blocked head (BH) are structurally prevented from binding actin, while the free head (FH) is constrained in a conformation with reduced ATPase activity (13–15). The IHM is closely associated with the super-relaxed state (SRX), a biochemical state characterized by very low ATP turnover, which is thought to play an important role in regulating energy consumption in cardiac muscle (2, 16, 17).

Altering the equilibrium between the closed/inhibited (IHM) conformation and an open/active conformation, in which the heads do not interact with each other (Fig. 1B, C), causes either excessive activation or enhanced sequestration, and this is a key molecular mechanism underlying cardiomyopathies (18–21). Mutations in cardiac myosin are a common cause of inherited hypertrophic (HCM) and dilated cardiomyopathy (DCM) (22). HCM is characterized by left ventricular wall hypertrophy, decreased left ventricular chamber volume, abnormally increased ejection fraction, and compromised relaxation. In contrast, DCM is typified by a thinner ventricular wall, increased ventricular chamber volume, and reduced ejection fraction and contractility (21, 23). Studies suggest that many of these myosin mutations affect the stability of the IHM and the balance between the closed/SRX and active/open states by perturbing interface interactions that stabilize the folded IHM conformation (18, 20, 24, 25). Mutations that disrupt the IHM are typically associated with the enhanced contractility of HCM, while mutations that stabilize the IHM are generally associated with reduced contractility (DCM), consistent with increased sequestration of myosin heads (24, 26). The DCM E525K mutation, located in the lower 50 kDa domain of the myosin head, has been studied extensively using biochemical and biophysical approaches. These studies reveal a puzzling contradiction in myosin activity, in which two-headed E525K myosin constructs exhibit an increased population of myosin in the autoinhibited IHM/SRX state (Fig. 1B), leading to altered force production, while isolated myosin heads (S1) demonstrate that E525K increases actin-activated ATPase activity, accelerates phosphate release, and enhances motility (26–29). The molecular basis for these opposite effects remains unclear. Our previous cryo-EM study of a wild-type IHM construct containing 25 heptads (250 Å) of tail revealed that cardiac myosin exists as a dynamic ensemble with two major IHM conformations, distinguished by the position of the S2 segment. In one state, S2 bends towards and interacts with the FH, forming an S2–FH docked conformation, whereas in the other, S2 remains detached from the FH, producing an S2–FH undocked conformation (30). This suggests that conformational flexibility within the S2 and FH regions may contribute to IHM function. Another recent structural and computational study also highlighted the dynamic nature of the cardiac myosin IHM and showed that the E525K mutation stabilizes the IHM conformation (31). However, despite these observations, the molecular mechanisms by which the E525K mutation plays a dual, opposite role by differentially modulating the autoinhibited IHM state and the active isolated S1 state remain unclear, limiting our understanding of how the mutation contributes to altered cardiac myosin regulation and DCM pathogenesis.

Here, we use cryo-electron microscopy (cryo-EM) to determine the structures of both a two-headed human β-cardiac myosin construct, with 25 heptads of the coiled-coil tail, and a single isolated myosin head (S1), each carrying the E525K mutation, and compare them with the corresponding wild-type (WT) structural data (30). We observe that the E525K mutant predominantly adopts the closed (IHM) conformation, and that, within this conformation, S2 is primarily in the docked position: there is less conformational variability relative to WT, particularly in the proximal S2 region. Interestingly, the charge reversal mutation does not introduce additional interactions, but instead redistributes the existing contacts and enhances the overall electrostatic attraction at the BH – S2 interface, consistent with the reduced mobility of S2 and overall stabilization of the docked IHM conformation. In contrast, isolated S1 reveals local conformational changes in the activation-loop and SH3-like regions associated with mutation-induced disruption of the conserved E525 – K484 salt bridge interaction. To further examine these mutational effects, we performed complementary molecular dynamics simulations (MD simulations) of both the WT and E525K docked IHM states: E525K was seen to strengthen BH–S2 electrostatic interactions and reduce S2 conformational flexibility, stabilizing the docked IHM state.

Together, our results reveal how a DCM-associated charge reversal mutation can exert distinct, opposite effects on cardiac myosin in different structural contexts. While the E525K mutation stabilizes the autoinhibited IHM through enhanced BH – S2 electrostatic interactions, it simultaneously promotes localized structural changes associated with activation of the isolated S1 head.

## Results

### The E525K mutation stabilizes the closed (IHM) conformation, with S2 predominantly in the docked state

We used cryo-EM to investigate how the DCM-associated E525K mutation alters the structure of cardiac myosin in its IHM state. Following particle picking and 2D classification, we observed numerous well-defined class averages corresponding to particles in the closed (IHM) conformation (Fig. 2A). In contrast to our previous wild-type dataset (30), which contained both closed (IHM) and open conformations during 2D classification, the E525K dataset did not yield any well-resolved 2D classes corresponding to open states. Instead, the particles appeared conformationally homogeneous, consistent with stabilization of the closed (IHM) conformation.

**Fig 2.**
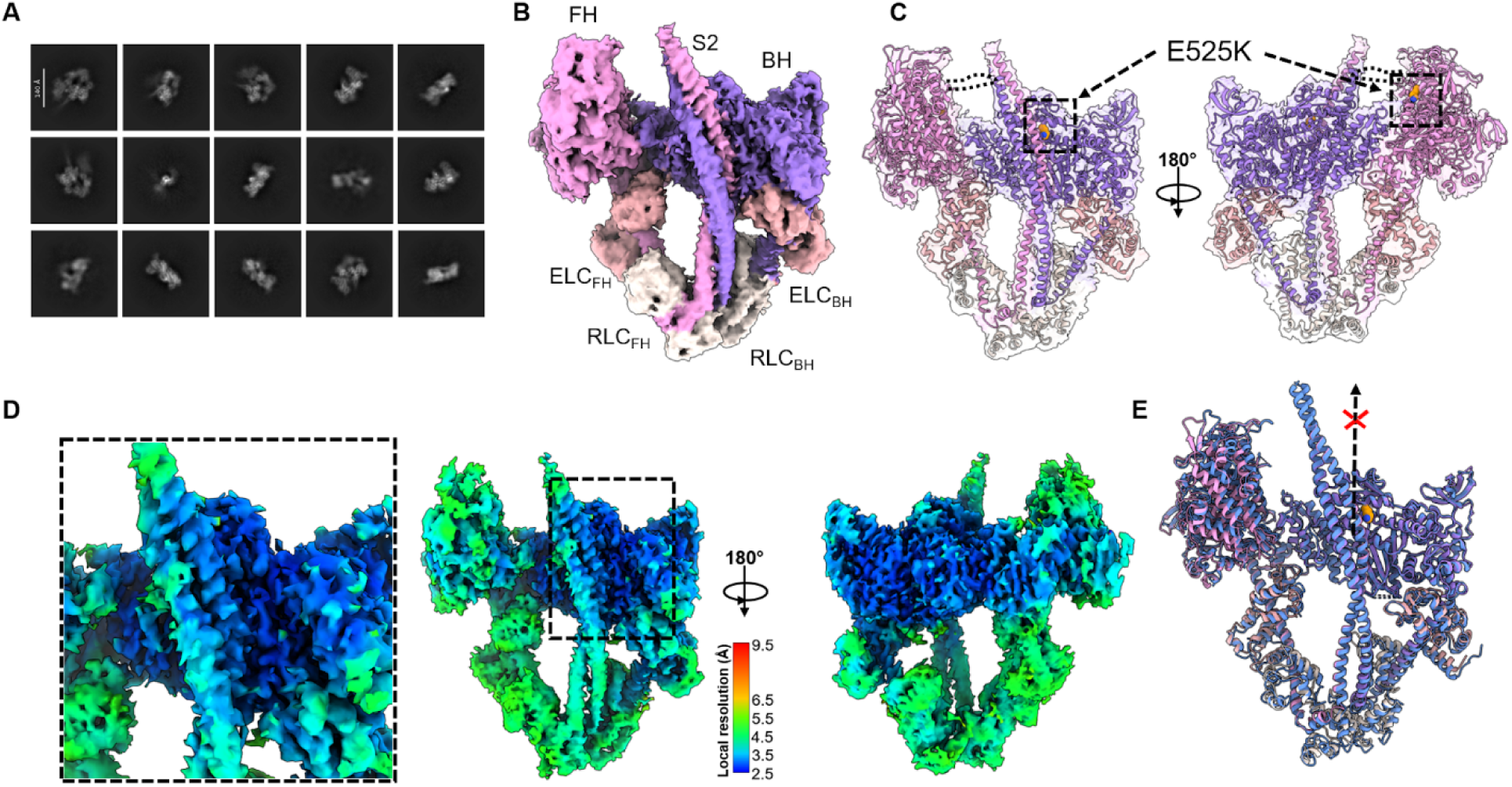
The E525K mutant stabilizes the closed IHM conformation. **A.** Representative 2D class averages of the E525K dataset showing well-defined particles only in the closed IHM conformation; different appearances are due to different IHM orientations. **B.** Cryo-EM reconstruction of the E525K IHM colored and labeled according to different subunits of IHM. **C.** Transparent cryo-EM density map with fitted atomic model, highlighting the location of residu K525 (orange) in both the BH and FH. FH Loop 2 and its potential interaction with S2 is illustrated as a dotted line between FH and S2. **D.** Local resolution map of the E525K reconstruction. The BH mesa – S2 interface is enlarged on the left to reveal higher local resolution, especially in the interacting S2 segment, indicating increased stability at this region in contrast to the less stable S2 in the WT (30). **E.** Structural alignment of WT and E525K IHM structures on BH. The E525K structure (pink) closely matches the docked WT conformation (blue), whereas the alternative WT undocked S2 conformation (dashed arrow with X (30)) i absent in the mutant, consistent with preferential stabilization of the docked IHM state.

Subsequent *ab initio* 3D reconstruction and refinement of the IHM particles produced a reconstruction at an overall resolution of 3.0 Å, as determined by the gold-standard FSC 0.143 criterion (Fig. S1). The density map confirmed, in three dimensions and at high-resolution, that the E525K mutant construct formed a typical IHM, with well-established head-head and head-tail interactions seen previously by cryo-EM in 15-heptad (14), 25-heptad (30) and 42-heptad (32) WT constructs. The resulting atomic model revealed that the mutant adopted an IHM conformation where the S2 and FH are arranged specifically in a docked configuration (Fig. 2B, C), with S2 residues ∼875 to 915 interacting with the BH mesa and bending towards the FH, where they can potentially interact with FH Loop 2 (Fig. 2C) (30). Structural alignment with the previously determined wild-type IHM structure showed that the mutant resembles the docked conformation observed in the WT; no alternative S2–FH undocked conformation was detected in 3D classification analysis of the mutant (Fig. 2E, S1). This suggests that stabilization of the closed over the open conformation is due to enhanced stability of the docked compared with the undocked conformation (30).

Local resolution analysis revealed variation across the mutant structure, with higher resolution of ∼2.5 Å observed at the BH mesa and proximal S2 interface relative to more peripheral regions (Fig. 2D). This signifies increased stability at the inter-domain interactions of the BH-mesa and S2. Notably, residue E525 is positioned on the BH mesa precisely at the interface with S2 (Fig. 2C, orange). Introducing an E K charge reversal at this site places a positively charged lysine in close proximity to Ring 1 acidic residues (residues: E894 – D906) within the interacting S2 segment (Fig. 2C).

### Conformational variability analysis reveals restricted S2 mobility in the E525K mutant

To assess the conformational heterogeneity within the interacting-heads motif, we performed 3D variability analysis (33) on both the E525K and existing WT datasets (30) and examined the extreme maps corresponding to frame 1 and frame 20 along the first principal component (PC1). PC1 accounts for the observed structural variability in the dataset and captures large-scale internal motions of the IHM. To enable quantitative comparison between the WT and mutant, atomic models were flexibly fitted into the low-resolution 3D variability maps (frame 1 and frame 20) using molecular dynamics flexible fitting (MDFF) (34).

In the wild-type reconstruction, PC1 captured the previously described transition between the S2–FH docked (Frame 1) and S2–FH undocked (Frame 20) conformations (Fig. 3 D, E). Quantitative analysis of these end states revealed angular displacements of ∼7.8° for the FH and ∼18.1° for the distal S2 segment, indicating coordinated motion in which both the FH and S2 move away from the FH–BH interface (Fig. 3F and Movie S1) (30). Consistent with this transition, the BH loop 2–Ring 1 and FH loop 2–Ring 2 interfaces are altered in frame 20, reflecting disengagement of S2 from the FH in the transition from the docked to the undocked conformation (Fig. 3D-F).

**Fig 3.**
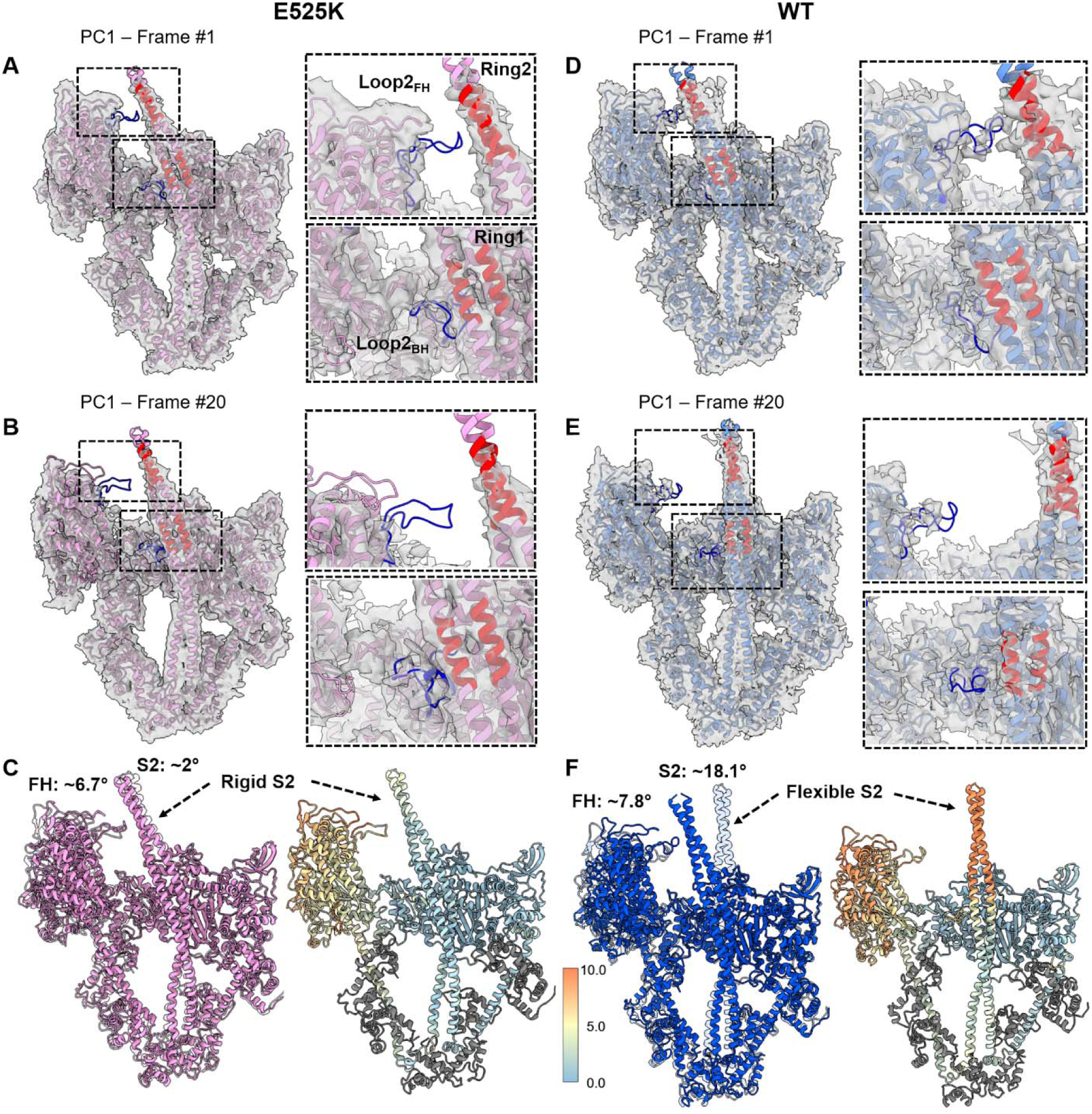
3D Conformational variability analysis reveals restricted S2 mobility in the E525K mutant. A, **B**. PC1 extreme conformations of E525K (frame 1 and frame 20) shown as transparent density maps with fitted atomic models. BH Loop 2 – S2 Ring 1 and FH Loop 2 – S2 Ring 2 interactions are highlighted in blue (Loop 2) and red (Ring 1, 2), with enlarged views shown alongside. **C,** Superposition of E525K frame 1 (pink) and frame 20 (transparent pink) aligned on the BH, highlighting limited movements. At right, frame 20 is color-coded by per-residue RMSD relative to frame 1 (blue-to-orange), showing conformational variability within the FH but little in S2. **D**, **E**, Corresponding PC1 extreme conformations of WT (frame 1 and frame 20). **F.** Superposition of WT frame 1 (blue) and frame 20 (transparent blue) aligned on the BH, revealing substantially larger S2 displacement than observed in E525K. The adjacent per-residue RMSD colored structure highlights increased conformational variability within the FH and distal S2 regions in WT.

In contrast, the E525K mutant exhibited markedly reduced structural variability along PC1. Comparison of the corresponding end maps shows both frame 1 and frame 20 adopting a docked IHM conformation in which S2 remains bent towards the FH (Fig. 3 A, B). Although the FH undergoes angular displacement of ∼6.7°, as observed in WT, the distal S2 segment displayed minimal motion, with an angular displacement of only 2° (Fig. 3C and Movie S2). Thus, while FH flexibility is largely retained in the mutant, S2 mobility is substantially suppressed. Close inspection of the BH loop 2–Ring 1 interface reveals that these contacts are largely preserved between the two end states, consistent with stabilization of the E525K IHM through electrostatic attraction at this interface (Fig. 3A, B).

### Enhanced and reorganized electrostatic interactions at the S2 – BH mesa interface direct S2 stabilization in the E525K mutant

To further investigate how the E525K substitution alters interfacial interactions, we analyzed the electrostatic contacts at the S2 – BH mesa interface. The bend in the S2 coiled-coil is located within residues ∼875–905, where S2 overlaps extensively with the blocked head (30). Within this region, Ring 1 from the FH coiled-coil helix, spanning residues 894–906, forms the primary interaction interface with the BH (Fig. 4A, E). Ring 1 is composed of multiple acidic residues, including E894, D896, D900, E902, E903, and D906, of which D896, D900, and E903 are oriented towards the BH mesa and are well positioned to mediate electrostatic interactions with some of its basic residues (35) (Fig. 4B, F). In the WT docked conformation, E525 forms a close interaction with K484 (∼3.0 Å), consistent with a stabilizing salt bridge on the BH surface (28). This interaction is abolished in the mutant due to the charge reversal at position 525, eliminating favorable contact with K484 (Fig. 4C, D and G, H). However, this loss is accompanied by a redistribution of electrostatic interactions within the S2 interface region, where the introduced lysine enhances the positively charged environment at the BH mesa and promotes stronger electrostatic complementarity with the acidic Ring 1 region of S2. In WT, within Ring 1, D900_FH_ interacts with both K526_BH_ and R652_BH_, with distances of ∼5.0 Å to each (Fig. 4A-D). In the E525K mutant, D900_FH_ instead interacts directly with the mutant lysine at position 525_BH_ (∼4.0 Å), while maintaining interaction with R652 (∼5.2 Å), resulting in a more concentrated electrostatic interaction network centered on residue 525 (Fig. 4E–H). In general, adjacent positively charged residues such as K525 and K526 are unlikely to simultaneously interact with the partner residues in the same direction due to electrostatic repulsion and steric constraints. Consistent with this, our analysis indicates a redistribution rather than an increase in interfacial contacts where K525 pushes K526 away and interacts with D900.

**Fig 4.**
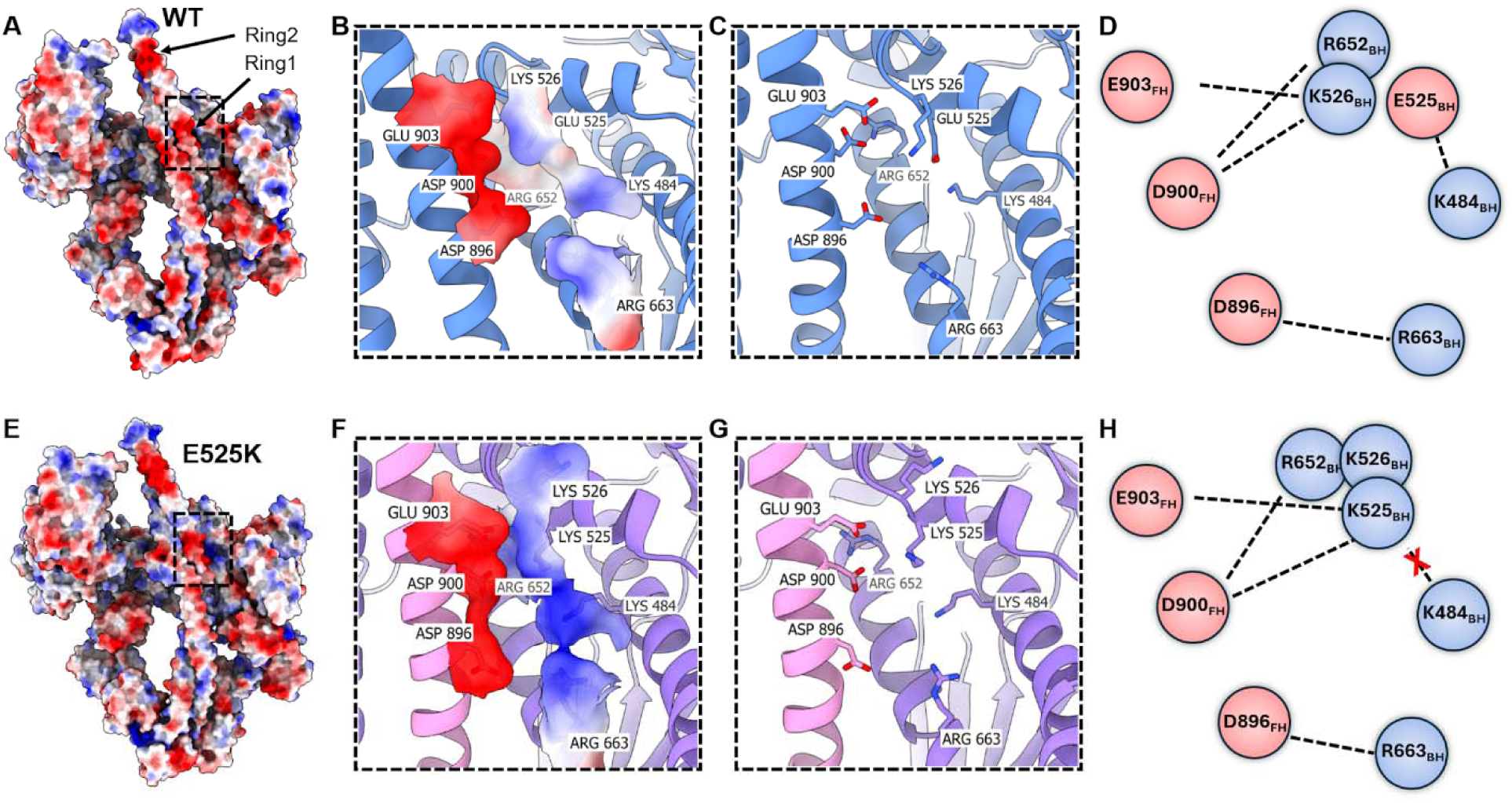
Enhanced electrostatic interactions at the BH mesa – S2 interface stabilize the E525K IHM. (**A–D**) WT and (**E–H**) E525K structures. **A, E.** Electrostatic surface potential maps showing the BH mesa and interacting S2 region, with the mutation site highlighted in a dotted box; blue, positive; red, negative. **B, F.** Enlarged views of the BH mesa – S2 interface. Electrostatic potential is mapped onto interface residues, illustrating the local charge environment surrounding residue 525 and the Ring 1 region of S2. In WT, E525 partially neutralizes the positively charged BH mesa, whereas the E525K substitution generates a more strongly positive electrostatic surface. **C, G.** Atomic models of the BH mesa – S2 interface showing residues involved in interfacial salt-bridge interactions. Key interacting residues are displayed as sticks and labeled accordingly. **D, H.** Schematic interaction network diagram of the interface with residues represented as nodes and electrostatic interactions as connecting lines. Positively charged residues are colored blue and negatively charged residues red. The E525K mutation breaks the E525 – K484 salt bridge and reorganizes the interaction network at BH – S2 interface by replacing E903-K526 and D900-K526 interactions with two newer ones involving K525 (E903-K525 and D900-K525), and enhancing the overall electrostatic complementarity between the BH mesa and the acidic Ring 1 region of S2.

In the WT, the presence of E525 partially offsets nearby positively charged residues (K484, K526, and R652), resulting in an evenly distributed electrostatic environment at the BH mesa (Fig. 4B). In contrast, substitution with lysine removes this local charge compensation and generates a more strongly positive electrostatic patch (Fig. 4F). This amplification of positive potential provides a more favorable environment for interacting with negatively charged residues within Ring 1 of S2, particularly D896, D900, and E903. This charge reversal mutation effectively concentrates electrostatic coupling at the BH–S2 interface, consistent with the reduced mobility of the S2 segment observed during 3D variability analysis, where distal S2 movement is markedly suppressed relative to wild type (Fig. 3). Together, these observations support a model in which the E525K substitution strengthens electrostatic complementarity at the BH–S2 interface by reorganizing and focusing positive charge electrostatic interactions within the BH mesa region. This enhanced electrostatic coupling stabilizes anchoring of the bent (docked) S2 coiled-coil and thereby constrains S2 dynamics. This phenomenon reduces the overall conformational heterogeneity compared with WT, and promotes stabilization of the closed, autoinhibited IHM conformation.

### Molecular dynamics simulations support stabilization of the docked IHM state by the E525K mutation

To investigate the impact of the E525K mutation on the overall dynamics of S2, we performed all-atom MD simulations of the docked WT and E525K IHM structures (Fig. 5A). S2 motion was analyzed by fitting a Bezier curve to the α-carbon center of mass of each pair of residues in the two α-helices of S2 (residues 843 to 957). An example for frame one of each simulation can be seen in Figure 5A, and an overlay of all 500 ns of simulation (one frame per ns) shows the relative movement of the S2 tail for each simulation (Fig. 5B). Although the curvature and movement of the S2 regions were highly variable, the E525K trajectory exhibited greater straightness and reduced maximum lateral deviation compared to WT (Fig. 5C and Movies S3 and S4). Consistent with E525K stabilizing the position of S2, we observed a decrease in the root mean square fluctuation (RMSF) of S2 in the mutant simulation compared to WT (Fig. 5D). Although both WT and mutant trajectories remained within the docked conformational ensemble during the 500 ns simulation time scale, the E525K mutant sampled a narrower range of S2 conformations than WT, consistent with a more constrained IHM geometry. This behavior is in excellent agreement with the cryo-EM 3D variability analysis, which showed reduced S2 mobility and preferential stabilization of the docked IHM conformation in the mutant (Movies S2 and S4).

**Fig. 5.**
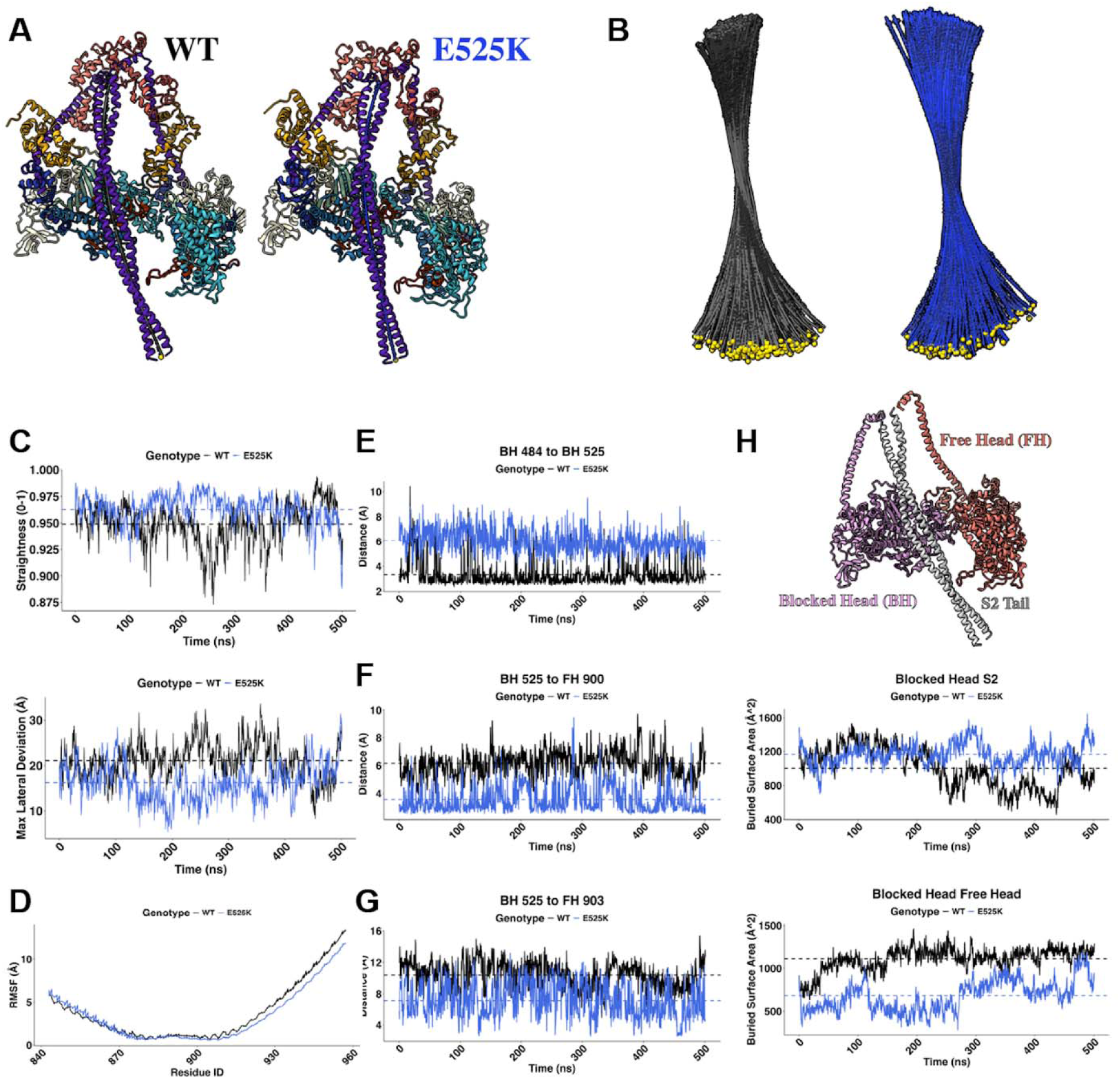
Molecular dynamics simulations of WT and E525K IHM cryo-EM structures reveal the dynamics of IHM stabilization by the E525K mutation. **A**. Representative WT and E525K IHM structures used for MD simulations. S2 flexibility was quantified by fitting a Bézier curve to the averaged α-carbon positions of the S2 coiled-coil. **B**. Overlay of Bézier curves from all simulation frames (one frame per ns), aligned using the S1 region of both myosin heads, illustrating the conformational space sampled by the S2 tail during the simulations in WT (black) and E525K (blue). **C**. Quantification of S2 geometry showing increased straightness and reduced maximum lateral deviation in E525K relative to WT, consistent with reduced S2 flexibility. **D**. RMSF analysis of the S2 tail after alignment to the central S2 region, showing reduced flexibility in E525K compared with WT. **E–G**. Charged side-chain distances between K484–K525, K525– D900, and K525–E903, respectively, demonstrate loss of the K484–K525 interaction and formation of closer interactions between K525 and the negatively charged S2 residues D900 and E903 in E525K. **H**. Buried surface area (BSA) analysis showing increased BH–S2 interface area after about 200 ns and a persistent decrease in BH–FH interface in the E525K simulation relative to WT.

We next examined residue pairs that form the principal interaction network surrounding mutant residue 525 in the BH mesa–S2 interface identified by cryo-EM (Fig. 5E-G, Fig. S2, S3). Consistent with the experimental cryo-EM structures, introduction of the positively charged lysine at position 525 both reorganized and enhanced the local electrostatic interactions. The average K525–K484 distance increased in E525K, consistent with charge-reversal loss of the local interaction present in WT (3.34 Å WT and 6.07 Å E525K; Fig. 5E). In contrast, distances between K525 and negatively charged residues within S2 Ring 1_FH_ decreased, including K525–D900 (6.16 Å in WT and 3.58 Å in E525K; Fig. 5F, S2) and K525–E903 (10.4 Å in WT and 7.14 Å in E525K; Fig. 5G), as charge attraction in the mutant replaced charge repulsion in the WT.

While the above Ring 1 residues of the FH interact with the BH mesa, the corresponding Ring 1 residues of the BH interact with BH Loop 2 (Fig. 3). We therefore examined whether the E525K mutation also alters the Loop 2_BH_–Ring 1_BH_ interaction network. Residue contact analysis revealed that, while Loop 2_BH_ contacts extended beyond the Ring 1 residues in WT, contacts in the E525K mutant were more concentrated within the Ring 1 region (Fig. S4). Together, these results indicate a coordinated reorganization and strengthening of the electrostatic interaction network at the BH–S2 interface (Fig. 5A-G, Figs. S2-S4) in agreement with the cryo-EM observations (Fig. 4D, H).

To more closely examine the interface between the heads and S2, we next quantified the buried surface area (BSA) across simulation time. These results showed an increase in average BSA between the BH and S2 in the E525K simulation compared with WT (1003 Å^2^ WT and 1164 Å^2^ E525K, Fig. 5H). Notably, this change was observed mostly in the latter half of the simulation after the structure was given time to adjust from the cryo-EM conformation. In addition to the BH–S2 interface, we also quantified the BH–FH buried surface area to examine whether stabilization of the S2 interaction alters inter-head packing. Although both simulations began with comparable BH–FH interfaces, the trajectories exhibited distinct interfacial behavior. While the WT simulation sampled a larger average BH–FH interface (∼1100 Å²), the E525K simulation maintained a smaller average BH–FH interface (∼650 Å²) throughout the trajectory (Fig. 5H). Thus, despite the overall stabilization of the docked IHM conformation by E525K, the mutant does not show increased stabilization at the BH–FH interface in the MD simulation. Instead, these results suggest that stabilization of the docked state can be driven primarily by the strengthened BH–S2 interaction, without requiring a larger BH–FH BSA. The reduced BH–FH BSA in E525K further indicates that the mutation may constrain the overall IHM architecture by preferentially stabilizing the BH–S2 interface rather than uniformly strengthening all interfaces within the IHM (Fig. 5H, S3, and Movies S3 and S4).

Although E525K is present in both myosin heads, its most direct structural effect is observed in the BH, where residue 525 participates in the BH–S2 interface. In the FH, residue 525 does not form a direct interaction with S2, and the FH remained highly flexible in the mutant, similar to WT. Nevertheless, MD simulations revealed changes in FH-associated interactions, including reduced BH–FH buried surface area, discussed above (Fig. 5H, S5), and a redistribution of FH Loop 2 interactions in E525K. To identify the FH loop 2 contacts that differed substantially between E525K and WT, we used a 40 percentage point difference in contact occupancies as a cut-off. In the WT simulation, FH Loop 2 formed contacts with residues within the activation-loop region, including E535, E536, C538, and M539, through K639 and K640. These interactions were not observed in the E525K simulation, whereas the mutant exhibited additional contacts with residues in the U50 region (Fig. S6). Thus, E525K in FH redistributes the interaction network of FH loop 2 more towards the U50 regions of the motor domain.

Together, these MD simulations demonstrate that E525K stabilizes the docked IHM state through enhanced BH mesa–S2 electrostatic interactions and reduced conformational freedom of the S2 coiled-coil.

### Structural changes in the isolated myosin head with E525K explain increased ATPase activity

To investigate whether the E525K mutation induces any structural changes within the isolated myosin head (i.e. independent of IHM formation), we determined cryo-EM structures of WT and E525K S1 and compared the atomic models (Fig. S7). Overall, both structures adopted the canonical pre-power stroke-like myosin conformation with highly similar global structures. However, local structural comparisons revealed significant conformational differences in three important regions of E525K S1 that were not observed in WT S1: the activation loop/H-loop and Loop 3 region (residues 526–593), the N-terminal SH3-like domain (residues 30–80) and the N-terminal region of the ELC (residues 43–125) (Fig. 6A-D and Movie S5).

**Fig 6.**
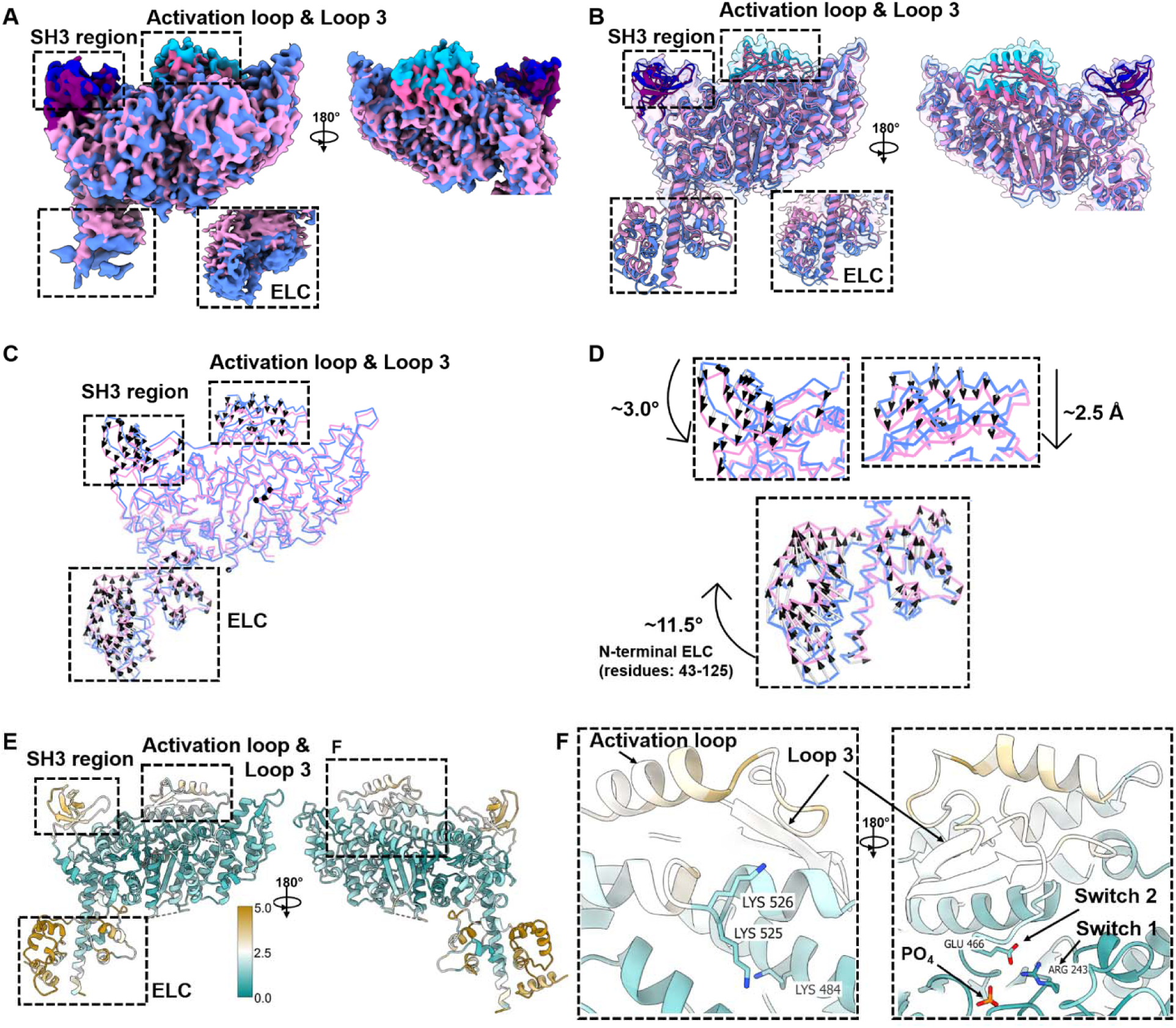
E525K induces local structural rearrangements of functional regions in the isolated myosin head. **A.** Cryo-EM density maps of WT (blue) and E525K (pink) S1 structures. Dotted boxes highlight regions that exhibit structural differences, including the activation-loop/Loop 3 region, N-terminal SH3-like domain, and essential light chain (ELC). The ELC shown here I derived from mouse skeletal muscle (see methods). The activation-loop/Loop 3 and SH3-like regions are highlighted in cyan and dark blue for WT, and magenta and purple for E525K, respectively. **B.** Corresponding transparent density maps with fitted atomic models. **C.** Visualization of mutation-induced structural rearrangements between WT and E525K S1 structures using displacement vectors with black arrow heads indicating the direction of the motion in the mutant structure compared to WT. **D**. Zoom in views of the major regions undergoing conformational changes. The arrows indicate the direction and magnitude of structural change. **E.** E525K S1 colored by per-residue RMSD relative to WT S1 (green to brown), highlighting localized deviations in the activation-loop/Loop 3 region, SH3-like domain, and ELC. **F.** Enlarged views of the E525K mutation site showing disruption of the E525–K484 interaction and associated rearrangements in the activation-loop and Loop 3 regions. The 180° rotated view at right highlights the adjacent Switch 1 and Switch 2 regions, including residues R243 and E466 that are associated with phosphate-release pathway.

In WT S1, residue E525 forms a conserved electrostatic interaction with K484 located near the relay helix region. In contrast, in the E525K mutant, this interaction is disrupted due to charge reversal at position 525, resulting in the observed local rearrangements within the activation-loop & Loop 3 regions (Fig. 6A-D). Quantitative structural analysis revealed that the center of mass of the activation-loop region shifts by 2.5 Å towards the nucleotide-binding region in the E525K structure (Movie S5). Importantly, both the activation-loop and Loop 3 contribute to the actin-binding interface of the motor domain. Loop 3 is positioned next to the switch 2 region and the conformational changes observed in the mutant extend beyond the immediate mutation site towards this neighboring switch 2, suggesting broader conformational coupling within the motor domain (Fig. 6F). Interestingly, both switch 1 and switch 2 contain key residues implicated in phosphate release through the proposed “back-door” pathway, including R243 in switch 1 and E466 in switch 2 (36) (Fig. 6F). These observations are in agreement with previous biochemical and simulation studies demonstrating that disruption of the E525–K484 interaction is associated with enhanced actin-activated ATPase activity, accelerated phosphate-release kinetics, and increased motility of the isolated S1 motor domain (28, 37). The structural changes observed here therefore provide direct evidence for intrinsic conformational changes within actin-binding regions of the motor domain of the E525K mutant that may contribute to increased actin-dependent motor activity.

Interestingly, additional conformational changes were also observed within the N-terminal SH3-like domain. Structural alignment revealed a shift/repositioning of the SH3 region in E525K S1, which is accompanied by an upward rotation of the ELC towards the motor domain compared to WT (Fig. 6A-D). Quantitative analysis showed that the SH3-like domain undergoes ∼3.0° rotation towards the N-terminal region of the ELC, while the N-terminal region of the ELC (residues 43–125) rotates ∼11.5° towards the SH3-like domain (Fig. 6D and Movie S5). The SH3-like domain has previously been shown to interact with the proline-rich N-terminal extension of the cardiac ELC, suggesting structural coupling between these two regions during the ATPase cycle (38). The coupled rearrangement of the SH3-like domain and ELC observed here suggests that the effects of the E525K mutation extend beyond the immediate activation-loop region and induce broader structural changes linking the motor domain and lever-arm regions (Movie S5). The results are consistent with an accelerated rotation of the lever arm observed in previous studies with the E525K mutant (28).

### E525K induces distinct conformational changes in isolated myosin heads not seen in the IHM

To determine whether the mutation-induced conformational changes observed in isolated S1 also occur in the heads of the IHM, we compared the free heads of the WT and E525K IHM and similarly the blocked heads (Fig. 7A, B). In contrast to S1, the activation-loop/Loop 3 region, SH3-like domain, and ELC of the FH and BH remained highly similar when comparing corresponding heads of WT and E525K IHMs. Although the E525–K484 salt bridge is disrupted by the E525K mutation in both the isolated S1 and IHM states, the structural consequences of this disruption vary strikingly between the two states (Figs. 4, 6). Whereas loss of this interaction in isolated S1 is associated with conformational rearrangements in the motor domain and ELC, these changes are not observed in either head within the IHM. This suggests that the extensive head-head and head-tail interactions present within the IHM constrain conformational flexibility of the motor domains and suppress structural changes that are otherwise present in isolated heads. Comparable constraints are also observed in the RLCs of IHM heads compared with S1. The RLC in S1 is not visible in the reconstruction (Fig. 6), suggesting flexibility around the RLC-ELC junction in the lever arm; this flexibility is minimal in the IHM, where the RLC is clearly resolved in both FH and BH (Fig. 3).

**Fig 7.**
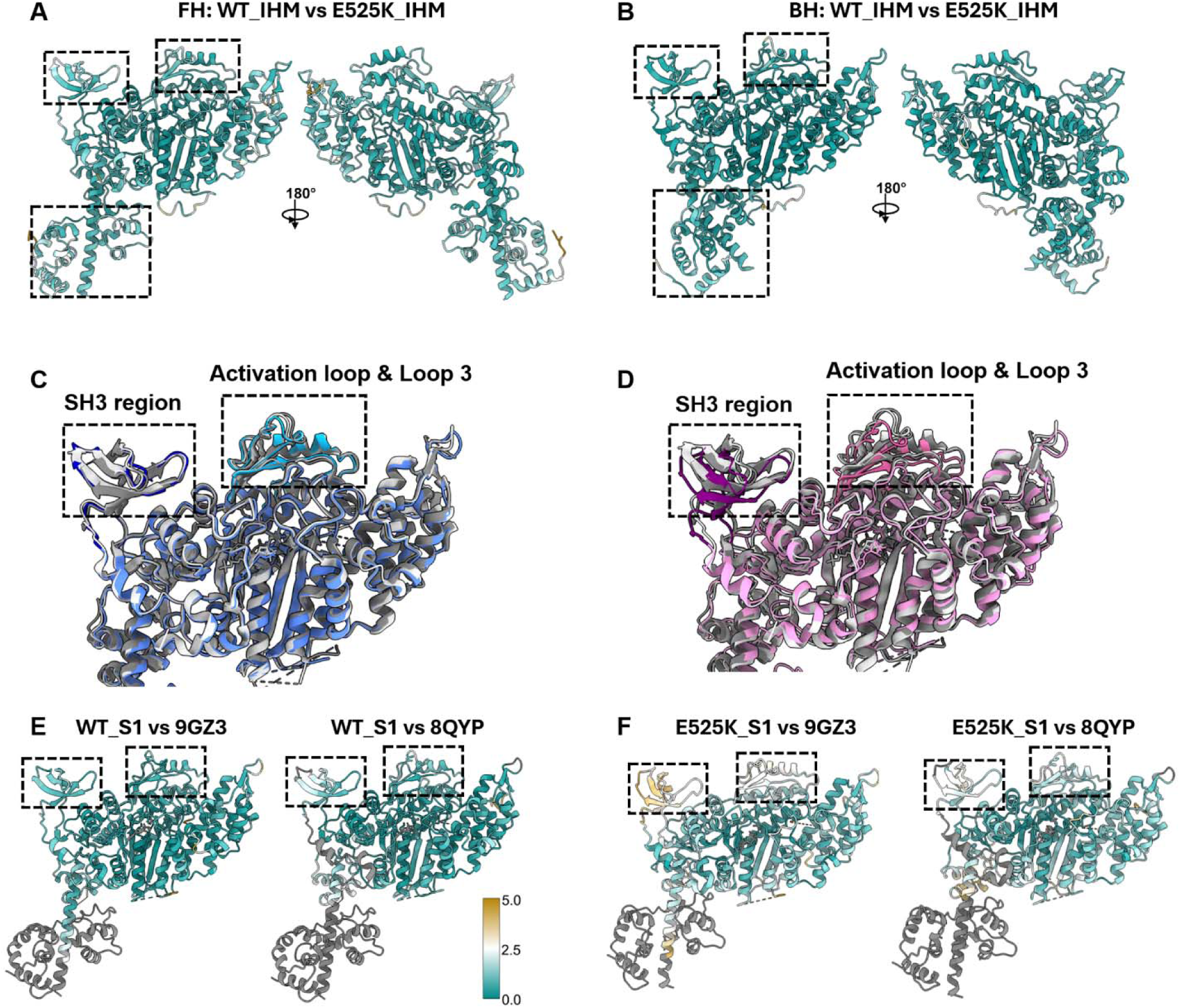
E525K-induced conformational changes within the isolated motor domain are absent in the IHM state and are distinct from intrinsic WT variability. A, **B.** Structural comparisons of WT and E525K FH and BH within the IHM. Mutant structures were color-coded by per-residue root mean square deviation (RMSD) relative to WT, mapped from low (green) to high (brown) deviation (see color bar on E). In contrast to isolated S1 (Fig. 6E), these regions show no detectable structural change in the IHM context. **C, D.** Structural comparison of previously reported myosin motor domain structures (9GZ3, human; light gray and 8QYP, bovine; dark gray) with our cryo-EM structures of WT S1 (blue, **C**) and E525K S1 (pink, **D**). **E.** WT and **F.** E525K structures were colored by per-residue RMSD relative to the reported apo motor domain structures (green to brown), further showcasing the intrinsic structural changes within the E525K motor domain compared to all WT structures. The regions that are unavailable to compare in previously reported structures are colored in gray.

To confirm that the conformational rearrangements observed in E525K represent intrinsic effects of the mutation rather than normal structural variability among myosin S1s, we compared our WT S1 structure with previously reported WT myosin motor-domain structures (PDB IDs: 9GZ3(32) (human) and 8QYP(36) (bovine)). Structural alignment revealed that the activation-loop/Loop 3 and SH3-like regions are highly conserved among WT motor domain structures despite differences in construct origin and experimental conditions (Fig. 7C, E). In contrast, the E525K S1 structure displayed distinct structural rearrangements within these regions compared with all the WT structures analyzed (Fig. 7D, F). Although ELC coordinates were unavailable in the previously reported structures, the conserved positioning of the WT motor domains in the regions available for comparison further supports that the conformational changes observed in E525K represent mutation-associated changes rather than inherent structural variability of the WT motor domain.

## Discussion

### Electrostatic effects and conformational dynamics of cardiac IHM

Electrostatic interactions have been proposed to play a central role in stabilizing the IHM, particularly through complementary charge distributions within different interfaces (20, 30, 39). Our findings are consistent with this view, suggesting that the E525K mutation not only reorganizes the interaction landscape at the BH–S2 interface but also strengthens overall electrostatic attraction in that region. Importantly, these electrostatic effects are closely tied to the conformational dynamics of the IHM, which we have recently demonstrated as a dynamic ensemble, with transitions between docked and undocked states contributing to regulation of myosin activity (30). In WT myosin, this is reflected in substantial mobility of the S2 segment and the coexistence of multiple IHM conformations (30). In contrast, stabilization of the BH–S2 interface in E525K is associated with reduced S2 flexibility while preserving FH movement, indicating that electrostatic stabilization can effectively restrict local motions without making the complex globally rigid. Notably, this region overlaps known HCM (E903G and D906G) and DCM (R904C) associated mutations, underscoring its functional importance in regulating head–tail interactions and IHM formation (24)

These observations are further supported by studies emphasizing the central role of charge balance at the BH mesa–S2 interface. Nag et al. (19) demonstrated that cardiomyopathy-associated mutations disrupting either positively charged residues on the mesa (R453C and R249Q) or negatively charged residues in proximal S2 (D906G) weaken motor domain binding to S2 and shift the equilibrium toward more open, active states. Similarly, Alamo et al. (20) showed that a large fraction of both HCM and DCM variants involved changes in charge, leading to distinct functional outcomes. In particular, mutations that increase negative charge in the S2 Ring 1 region (R904C) are predicted to strengthen BH–S2 interactions and stabilize the IHM, whereas mutations that reduce negative charge (D906G) weaken these interactions and favor open active states.

This coupling between electrostatics and dynamics suggests a mechanism in which mutations such as E525K stabilize the IHM by narrowing the accessible conformational landscape, thereby reducing the likelihood of transitions to more open, active states. More broadly, these observations show how subtle changes in charge distribution can tune the balance between structural stability and conformational flexibility in large macromolecular assemblies.

### E525K stabilizes the IHM against local environmental perturbations

The WT SRX/IHM structure is known to be sensitive to environmental conditions such as ionic strength, pH, and temperature, which can shift the equilibrium between closed IHM and open active conformations (26, 27, 39, 40). This intrinsic sensitivity reflects the dynamic nature of the BH–FH, BH–S2, and FH–S2 interfaces in the IHM, where electrostatic interactions play a central role in stabilizing the folded conformation (2, 20, 30), as discussed above. In contrast, the E525K charge-reversal mutation stabilizes the IHM, reducing this environmental susceptibility. We have measured the SRX and IHM populations in constructs of WT and E525K cardiac myosin across a range of ionic strengths (26, 27). In low salt (20 mM), both had a high % of IHM and SRX. At higher salt (150 mM), the E525K mutant maintained a high SRX/IHM population, whereas WT showed a dramatic decrease compared with low ionic strength (26, 27). This suggests that the mutation strengthens electrostatic interactions that are weakened in the WT under high salt conditions, consistent with our cryo-EM observations. Our findings also explain EM observations showing that conventional negative staining almost eliminates the apparent IHM population in WT but not in the E525K mutant, due to surface interactions with the carbon substrate, suggesting that the mutant is more resistant to environmental disruption (41). The enhanced stability of the IHM suggests a structural basis for the increased SRX population and reduced contractility associated with the mutation (26, 27).

### Contrasting structural effects of E525K in the IHM and isolated S1

In contrast to its stabilizing effect in the IHM, the same E525K mutation induces localized structural rearrangements in the isolated S1 motor domain that are not seen in the heads of the IHM. Previous biochemical studies have demonstrated that E525K S1 exhibits increased actin-activated ATPase activity, higher actin affinity, and faster motility compared to WT (27, 28). These effects have been attributed to disruption of a conserved intramolecular salt bridge between E525 and K484 near the relay helix, thereby increasing local flexibility and promoting transitions through the ATPase cycle (28). Our cryo-EM structures of S1 and the IHM provide direct structural evidence for this mechanism (Figs. 4-6). In addition, comparison of WT and E525K S1 reveals significant conformational changes in the activation-loop/Loop 3 region and in the N-terminal SH3-like domain, accompanied by repositioning of the ELC toward the motor domain (Fig. 6 and Movie S5). These structural perturbations potentially extend beyond the mutation site towards the neighboring switch 2 region, which is part of the phosphate-release pathway. In contrast, in the two-headed IHM configuration, the same mutation shows no structural changes in these regions (Fig. 7). Instead, the IHM is stabilized and S2 flexibility is reduced (Figs 3, 4; (31)), explaining stabilization of the autoinhibited state and the increase in SRX population (26, 27). Our structures and simulations further reveal the molecular basis of this stabilization, where the introduced lysine at position 525 reorganizes and strengthens electrostatic interactions at the BH – S2 interface through interaction with acidic residues within the S2 Ring 1 region. This strengthened interfacial coupling constrains S2 flexibility and reinforces the autoinhibited IHM state, limiting the number of isolated single heads available for actin interaction.

These findings suggest that the ultimate effect of E525K in cardiac muscle reflects the interplay between the two opposing molecular consequences: stabilization of the IHM (Figs 3, 4), which decreases the number of myosin heads available for interaction with actin, and enhanced motor activity of individual heads when they are released from the IHM and interact with actin. Because force production at the sarcomere level depends not only on the kinetics of individual motors but also on the number available for actin interaction, the increased sequestration of heads in the IHM appears to be the dominant effect in vivo, explaining the hypo-contractile DCM phenotype despite enhanced biochemical activity of isolated motor domains (28). Collectively, our findings support a model in which the structural effects of E525K are state-dependent, producing local conformational changes in isolated S1, while simultaneously stabilizing the autoinhibited IHM state of the two-headed molecule without significantly changing the conformation of the individual heads.

### Implications of E525K-mediated IHM stabilization on thick filament regulation and DCM

A recent cryo-EM structure of the human cardiac thick filament at 6.0 Å resolution has revealed that the IHM in the filament can adopt two distinct conformations, where the interacting motor domains (BH and FH) are either tilted or perpendicular to the filament axis (6). Comparison of our E525K cryo-EM map to the tilted and horizontal IHMs of the filament (EMDB ID: EMD-29722) reveals that the E525K IHM, characterized by a bent S2 docked onto the FH, aligns well with the tilted and less so with the horizontal IHM. In contrast, the conformational freedom observed in the WT construct, which exhibits both S2 – FH docked (PDB ID: 9YOP) and S2 – FH undocked (PDB ID: 9YRG) structures allows conformational similarity with both tilted and horizontal configurations (30). Thus, the intrinsic flexibility of the WT IHM enables sampling of multiple conformational states consistent with the native filament, whereas the E525K mutation restricts this conformational landscape.

This restriction has important implications for thick filament function and regulation. Stabilization of the BH – S2 interface in E525K likely promotes sequestration of myosin heads along the filament backbone. Rather than equilibrating between open and closed (IHM) states, the E525K mutation strengthens the BH – S2 interface and stabilizes the IHM conformation. This holds the two myosin heads more tightly within the folded-back IHM state, limiting their ability to transition into actin-engaged, force-generating conformations. Such a shift in conformational equilibrium would reduce the structural diversity of myosin states within the filament and constrain the flexibility required for efficient recruitment of myosin heads into the open state during muscle contraction (Fig 1B). Our findings provide an understanding of the pathogenesis of dilated cardiomyopathy caused by this single-point E525K variant. By pinpointing this mutation’s non-physiological stabilization of the IHM through strengthening of the BH-S2 interface, our results provide a clear biophysical mechanism linking a mutation-induced molecular-level effect to clinical ventricular heart remodeling that leads to a dilated heart exhibiting hypodynamic contraction, unaffected relaxation, and decreased metabolism.

## Materials and Methods

### Protein construction, expression, and purification

The human β-cardiac myosin 25-heptad HMM construct was expressed and purified as previously described (26, 27). Briefly, the myosin *MYH7* gene (GenBank: AAA51837.1) was engineered with a C-terminal arrangement of GCN4 leucine zipper (sequence: MKQLEDKVEELLSKNYHLENEVARLKKLVGER), a short linker (GSGKL), EGFP, Avi-tag (GLNDIFEAQKIEWHE), and FLAG-tag (DYKDDDDK), and cloned into the pDual shuttle vector. The S1 construct (amino acids 1–842) containing the motor domain and associated essential and regulatory light chains was generated in parallel using the same expression system but had only the C-terminal EGFP, Avi-, and FLAG-tags. The E525K mutation was introduced using Quikchange site-directed mutagenesis (Agilent Technologies), and all constructs were verified by DNA sequencing. Recombinant adenovirus was generated by Vector Biolabs at a titer of ∼10 plaque-forming units per mL (pfu/mL). Protein expression was carried out in differentiated C2C12 mouse skeletal muscle cells as described previously(26). The expressed human beta-cardiac myosin heavy chain constructs containing mouse skeletal essential and regulatory light chains from the C2C12 cells (42) were purified using FLAG affinity chromatography.

### Cryo-EM grid preparation and data collection

Purified myosin constructs were flash-frozen in liquid nitrogen with 5% sucrose solution and stored at-80 °C until further use. For grid preparation, the protein was diluted to ∼1.5 μM immediately prior to vitrification. The E525K HMM construct was diluted in buffer containing 10mM MOPS, 2mM MgCl2, 0.5mM ATP, pH 6.8, whereas the WT and E525K S1 constructs were diluted in the same buffer with 20mM KCl at pH 7.4. A volume of 3 μL was applied to a Quantifoil R1.2/1.3 UltraAuFoil 300 mesh gold grid, which had been glow-discharged at 25 mA for 30 seconds (EasiGlo, Ted Pella). Grids were blotted for 3 seconds under 100% humidity at 25 °C and vitrified in liquid ethane using a Vitrobot Mark IV. Cryo-EM data were collected on a Titan Krios transmission electron microscope (ThermoFisher Scientific) operated at 300 kV and equipped with a Gatan K3 direct electron detector in counting mode (Table S1). Separate datasets were collected for each construct under identical imaging conditions. A total of 8544, 4218 and 6319 movies were recorded for E525K HMM, WT S1, and E525K S1 respectively at a nominal magnification of 105,000×, corresponding to a calibrated pixel size of 0.83 Å. A total electron dose of ∼50 e /Å² was applied over 40 frames with an exposure time of 2.5 s. Data were collected over a defocus range of −1.0 to −2.0 μm using SerialEM (43).

### Cryo-EM data processing, model building and refinement

All the HMM and S1 datasets were processed independently using cryoSPARC software (44). Movie frames were motion-corrected using Patch Motion Correction, and contrast transfer function (CTF) parameters were estimated using Patch CTF Estimation. Particles were automatically picked using a blob picking approach, followed by Topaz particle picker (45) which had been pre-trained separately for the HMM and S1 datasets for enhanced particle picking. For E525K HMM, an initial dataset of ∼3.0 million particles was subjected to several rounds of 2D classification, yielding ∼150,000 high-quality IHM particles. These particles were further processed by 3D classification to separate any distinct conformational states. Particles from the best resolved classes were selected and refined using homogeneous refinement followed by non-uniform refinement in cryoSPARC, resulting in a final reconstruction at ∼3.0 Å resolution based on the gold-standard Fourier shell correlation (FSC) 0.143 criterion (Fig. S1). The map was subsequently post-processed with B-factor sharpening in cryoSPARC. The WT and E525K S1 image processing follows a similar workflow with several rounds of 2D classification, followed by ab initio reconstruction, heterogeneous refinement, homogeneous refinement, and non-uniform refinement in cryoSPARC to obtain the final S1 reconstructions (Fig. S7).

An initial atomic model of the E525K IHM was generated using the recently determined human WT β-cardiac myosin structure in the docked state (PDB ID: 9YOP (30)) as a starting model. Initial models for the WT and E525K S1 structures were generated by homology modeling using the bovine cardiac myosin S1 structure in the pre-powerstroke state (PDB ID: 5N69) as the template, with the heavy chain replaced by the human β-cardiac myosin sequence and the associated mouse skeletal essential light chain. All models were initially rigid-body fitted into the cryo-EM density using UCSF ChimeraX (46), followed by molecular dynamics flexible fitting (MDFF) implemented in Namdinator (47). Subsequently, iterative real-space refinement was performed in Phenix (48). Intermediate manual adjustments of side chains and loops were performed in Coot (49). Model quality was assessed using MolProbity (50). The final cryo-EM density maps and corresponding atomic coordinates have been deposited in the EMDB and the PDB under accession codes EMD-78337, 37OA (E525K IHM); EMD-78401, 37PY (WT S1); EMD-78403, 37QC (E525K S1) (Table S1).

### Molecular dynamics simulations

Structures determined by cryo-EM were prepared for MD simulations using the AMBER20 simulation package and the ff14SB force field. Water molecules were modeled using the TIP3P water model, whereas metal ions were parameterized using the Li and Merz parameter set. ADP and inorganic phosphate (Pi, modeled as H PO) were described using the GAFF2 force field. Partial charges for ADP and Pi were derived from a restrained electrostatic potential (RESP) fitting to quantum mechanical calculations performed with ORCA. Hydrogen atoms were added using the leap module in AMBER, and an N-methylamide (NME) cap was added to the C-terminus of each construct. Each protein was solvated in an explicit water box with a minimum buffer of 15 Å between the protein surface and the box boundary, and neutralizing counterions were added to maintain charge neutrality.

Each system undergoes a three-stage energy minimization protocol. First, hydrogen atoms were minimized for 1,000 steps while restraining all heavy atoms with a force constant of 100 kcal mol ¹ Å ². Second, solvent molecules were minimized for 1,000 steps with 25 kcal mol ¹ Å ² restraints applied to all protein atoms. Finally, all atoms were minimized for 8,000 steps, with 25 kcal mol ¹ Å ² restraints applied to all protein backbone atoms (N, Cα, C, and O). Following minimization, each system was gradually heated to 310 K through three successive 1 ns heating stages under the NVT ensemble while maintaining 25 kcal mol ¹ Å ² restraints on the backbone atoms. The systems were then equilibrated under the NPT ensemble in six successive stages. During the first four stages, backbone restraints were progressively reduced from 25 to 1 kcal mol ¹ Å ² over 5 ns per stage. The fifth stage consisted of 5 ns of unrestrained equilibration, followed by a final 50 ns unrestrained equilibration to ensure system stability before production simulations.

The SHAKE algorithm was applied to constrain hydrogen-atom bonds, and long-range electrostatic interactions were treated using the Particle Mesh Ewald (PME) method. Hydrogen mass repartitioning was employed to enable a 4 fs integration time step during heating, equilibration, and production simulations. A 10 Å cutoff was used for short-range nonbonded interactions. Independent simulations were performed for the WT and E525K IHM structures. Following equilibration, each system was subjected to 500 ns of production MD simulation under the NVT ensemble, yielding a total simulation time of 1 μs. The coordinates were saved every 10 ps for subsequent analysis.

### Molecular dynamics analyses

Structural flexibility of the S2 coiled-coil was quantified by fitting a Bézier curve to the averaged positions of paired Cα atoms from the two S2 helices at each simulation frame. From the fitted curves, straightness, and maximum lateral deviation parameters were calculated to compare S2 curvature between the WT and E525K trajectories. Straightness was calculated as the ratio of the end-to-end distance to the arc length of the fitted Bézier curve, with values near to 1 indicating a straighter S2 coiled-coil. Maximum lateral deviation was calculated as the maximum perpendicular distance of the Bézier curve from the straight line connecting its two endpoints which provides a measure of S2 bending.

The root-mean-square fluctuation of the S2 region was also calculated to assess local conformational flexibility. Buried surface area between the BH–S2, BH–FH, and FH–S2 interfaces was calculated throughout the simulations to monitor changes in inter-domain packing over time. Distances between selected residue pairs were calculated using CPPTRAJ. Intramolecular contacts were identified using the *nativecontacts* command in CPPTRAJ, with two residues considered to be in contact if any pair of atoms were within 5 Å. Contacts were retained for further analysis only if they were present for more than 10% of the simulation trajectory. Contacts were considered differentially populated between WT and E525K if their occupancies differed by more than 40 percentage points. Because each genotype was represented by a single simulation trajectory, statistical comparisons of contact occupancies were not performed.

## Acknowledgements

The authors thank C. Ouch, K. Song, and C. Xu (UMass Chan Medical School cryo-EM Facility) for cryo-EM training and imaging support, as well as K. H. Lee and G. Hendricks (UMass Chan Medical School Electron Microscopy Facility) for conventional EM training.

## Author contributions

R.G. and A.K.S. conducted cryo-EM experiments; prepared grids and collected cryo-EM datasets; processed, analyzed data and carried out reconstruction. R.G. performed preliminary model building. A.K.S. processed all cryo-EM data and built atomic models, performed structure analysis, and database deposition. K.Z.R. performed simulations and structural analysis. J.G. and C.M.Y. contributed to expression and purification of myosin samples. A.K.S. interpreted the data, prepared figures and wrote the manuscript with R.C. and R.P. All authors discussed the results and approved the final version of the manuscript. Funding acquisition, project administration, and supervision: A.K.S., C.M.Y., M.R., R.C., and R.P.

## Data availability

The cryo-EM density maps and corresponding atomic models have been deposited into EMDB and PDB under accession codes EMD-78337, 37OA (E525K IHM); EMD-78401, 37PY (WT S1); EMD-78403, 37QC (E525K S1). PDB data used to build the models were 9YOP and 5N69. In addition, the following cryo-EM map EMD-29722 and PDB models were used for comparisons: 9GZ3, and 8QYP.

