## Supplementary Information for "The dilated cardiomyopathy E525K mutation stabilizes the cardiac myosin interacting-heads motif while activating the isolated motor domain"

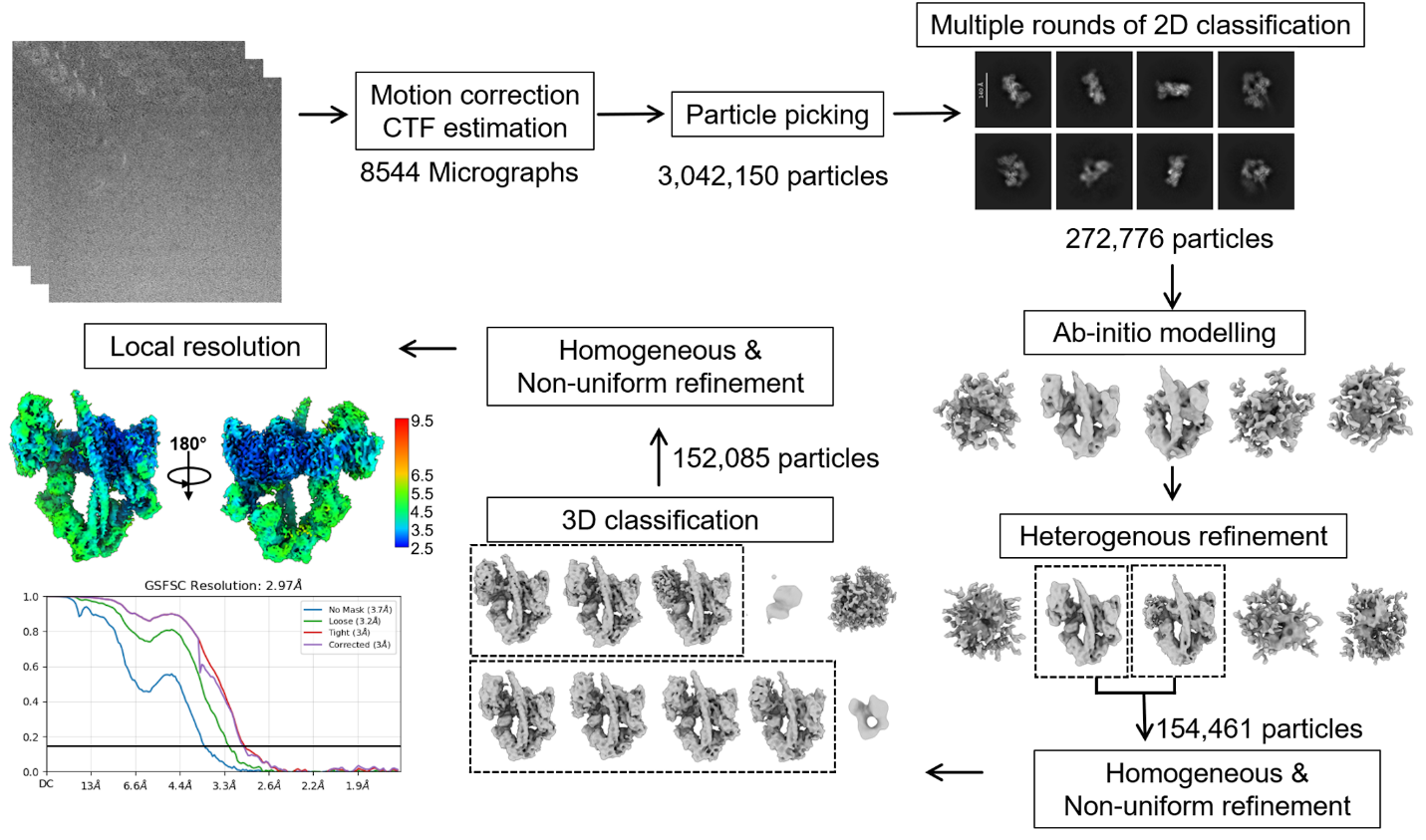

**Fig. S1. Cryo-EM data processing workflow of E525K-IHM.** Schematic overview of the image processing workflow for E525K-IHM, including motion correction, CTF estimation, blob and template-based particle picking, multiple rounds of 2D classification to separate junk particles from IHM, ab initio reconstruction and heterogeneous refinement for particle curation, and 3D classification leading to the final reconstruction, colored by local resolution.

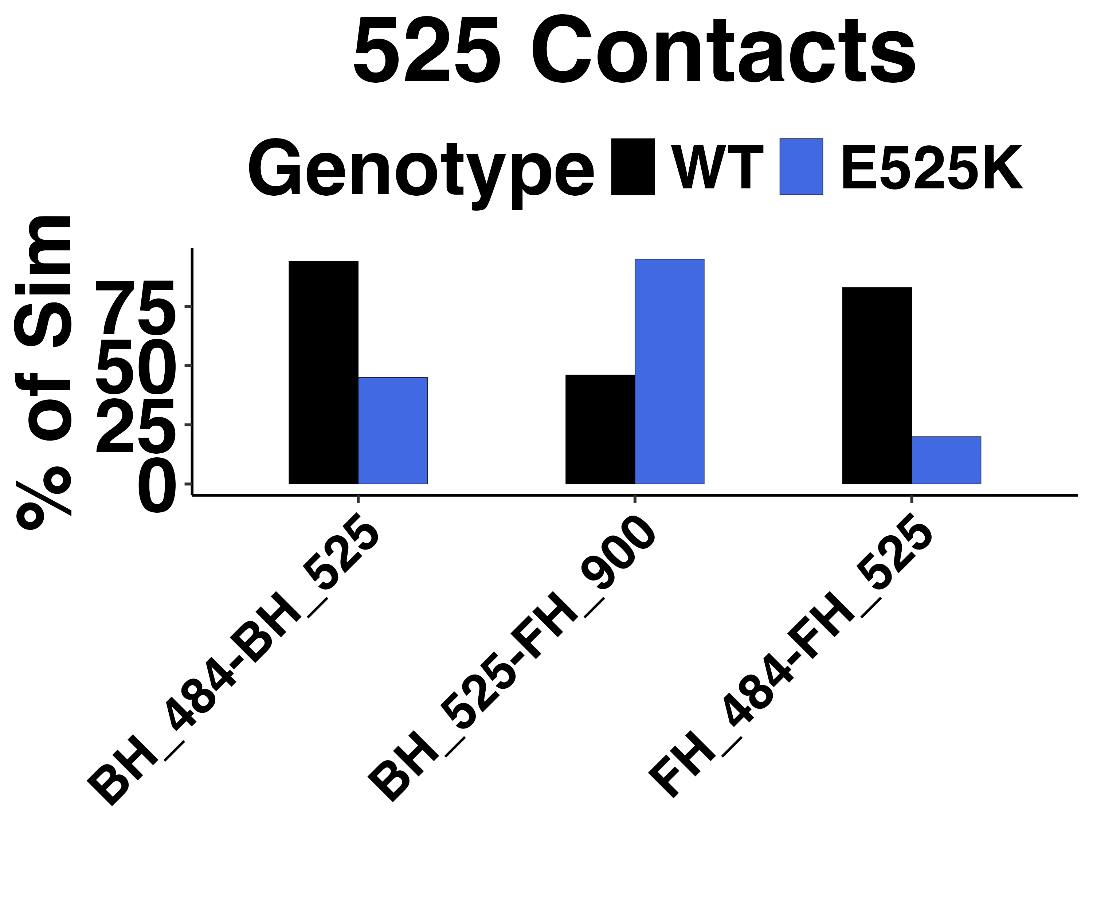

**Fig. S2. Contacts involving residue 525 during MD simulations.** Residue-contact analysis centered on position 525 that differed by more than 40 percentage points between WT and E525K. The E525K mutation abolishes the BH E525–K484 interaction while promoting formation of contacts between K525_BH_ and D900_FH_. The mutation also abolishes the FH E525–K484 interaction.

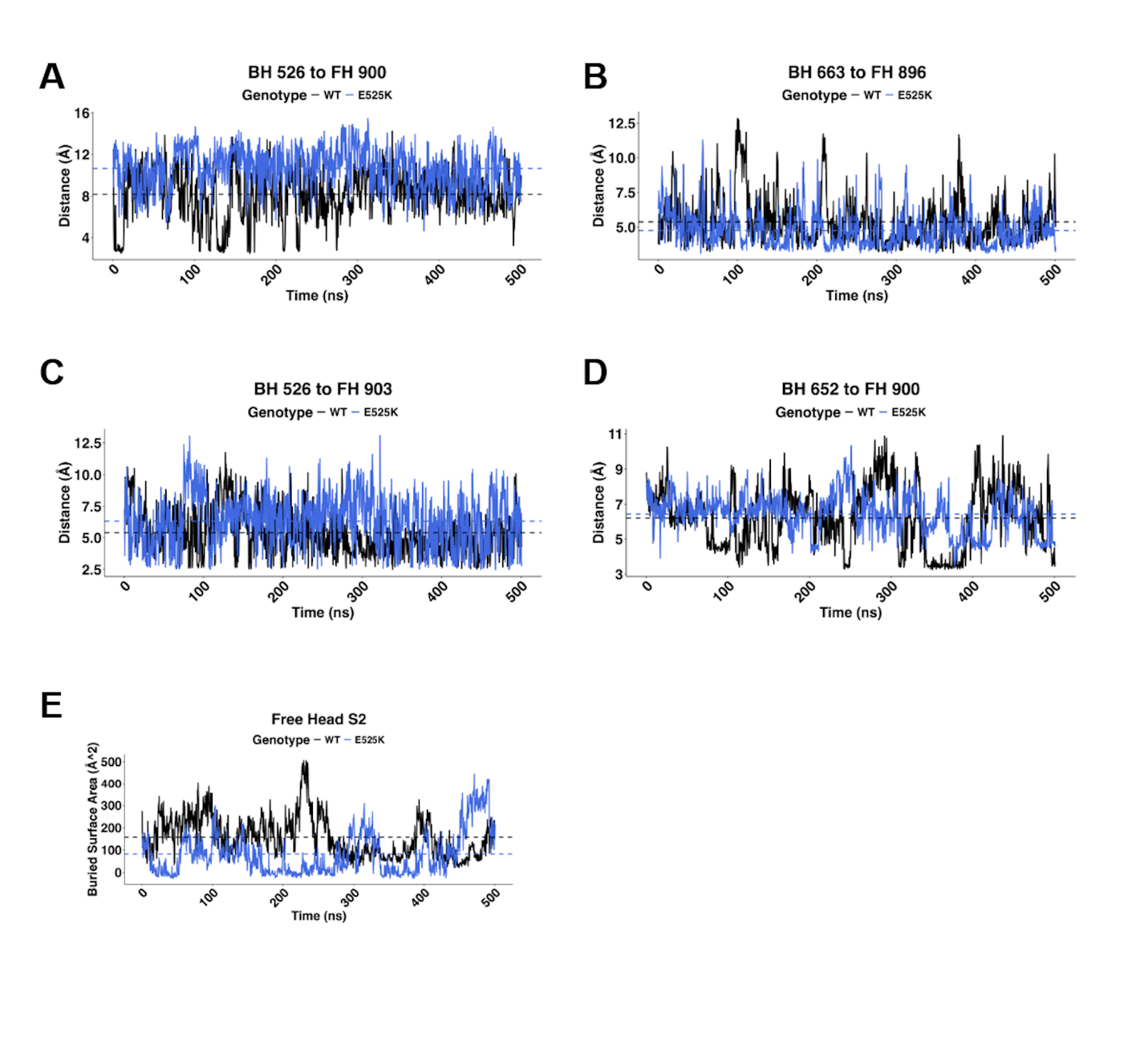

**Fig. S3. Distance and buried surface area analyses from MD simulations.** (A–D) Time-dependent charged side-chain distances for selected residue pairs at the BH–S2 interface: K526_BH_–D900_FH_ (A), R663_BH_–D896_FH_ (B), K526_BH_–E903_FH_ (C), and R652_BH_–D900_FH_ (D). (E) Buried surface area between the free head and S2 throughout the simulations. Together, these analyses show that the E525K mutation reorganizes interactions at the BH–S2 interface while altering the interdomain contact areas sampled during the MD simulations.

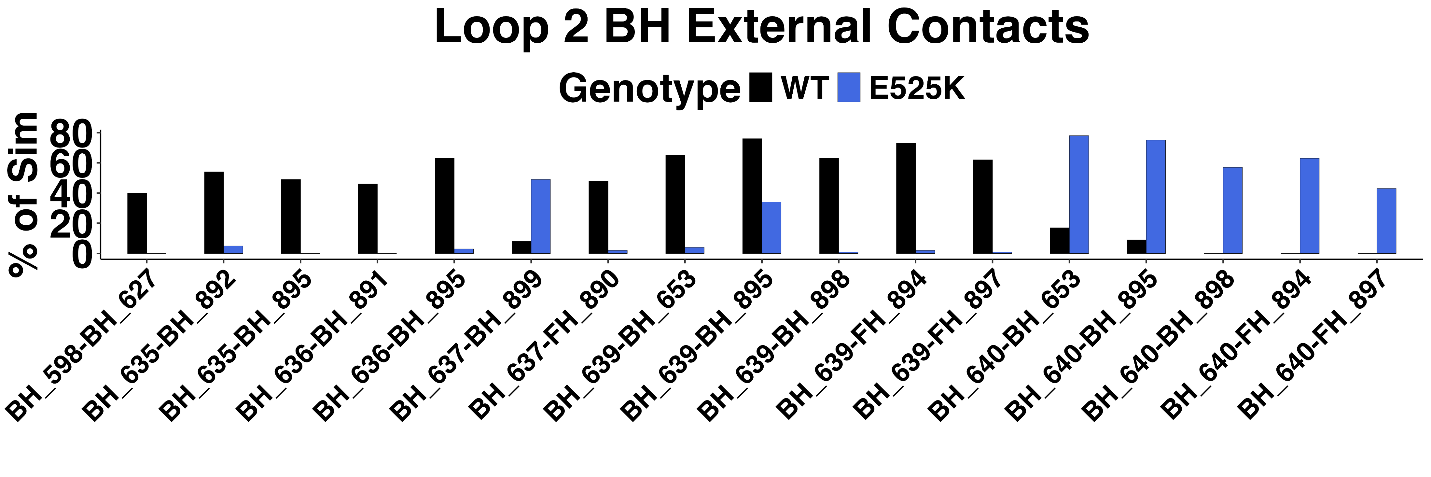

**Fig. S4. Contact network of BH Loop 2 during MD simulations.** Contacts between residues within BH Loop 2 (residues 622–645) and residues outside Loop 2 are shown. Only contacts whose occupancies differ by more than 40 percentage points between WT and E525K simulations are shown. This analysis shows that BH Loop 2 interactions in E525K are more concentrated within the corresponding Ring 1 region (residues 894–906) compared to interactions that extend beyond Ring 1 in WT.

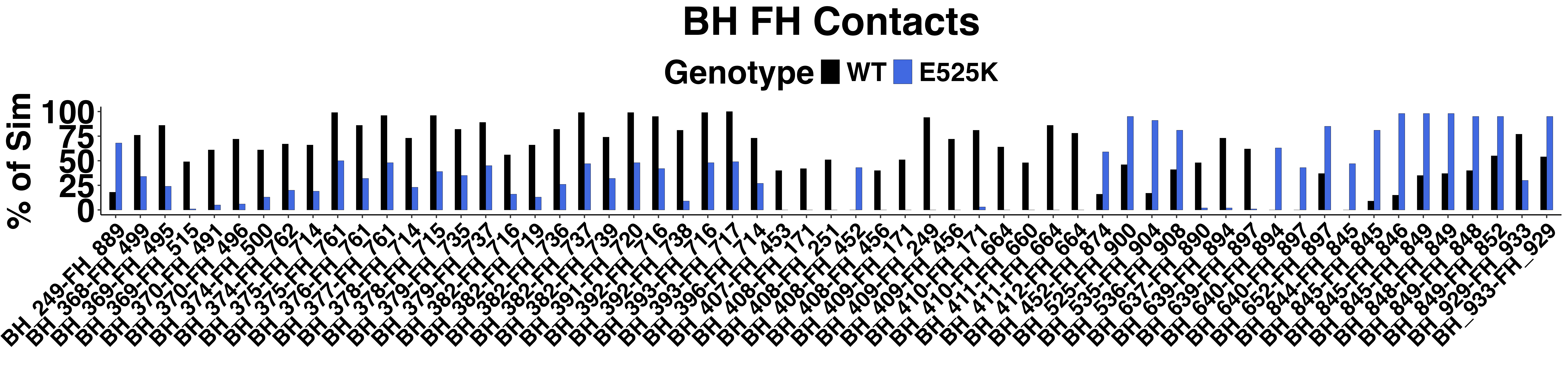

**Fig. S5. Inter-head contacts between the blocked head and free head during MD simulations.** Residue-contact analysis of the BH and FH interface. Contact occupancies are shown for WT and E525K simulations, illustrating differences in inter-head interactions sampled during the trajectories. Only contacts whose occupancies differ by more than 40 percentage points between WT and E525K simulations are shown. Overall, this analysis reveals substantial redistribution of BH–FH contacts in E525K relative to WT, consistent with the reduced average BH–FH buried surface area observed in the mutant simulation.

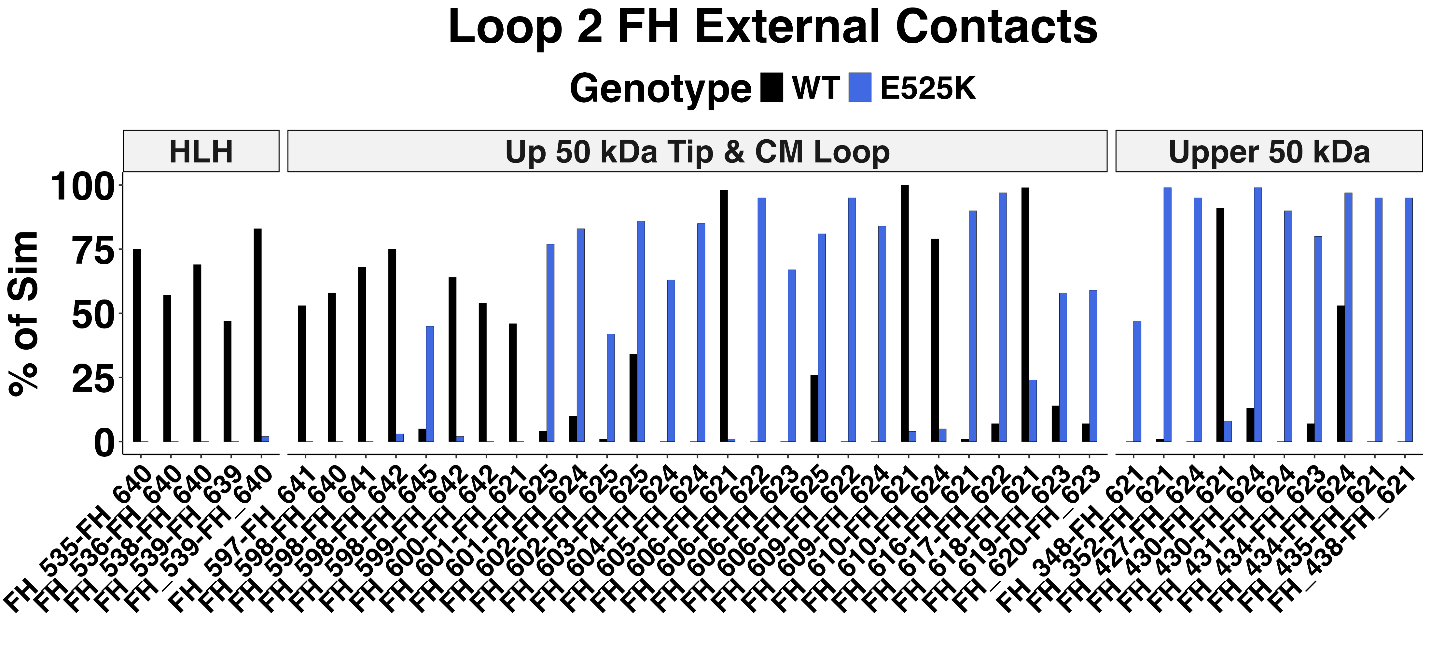

**Fig. S6.** **Contact network of FH Loop 2 during MD simulations.** Contacts formed between FH Loop 2 (residues 622–645) and residues outside the loop are shown. Contact partners are grouped according to their corresponding myosin subdomains. Only contacts whose occupancies differ by more than 40 percentage points between WT and E525K simulations are shown. This analysis shows that E525K redistributes the interaction network of FH Loop 2, with loss of contacts observed in the activation-loop region and increased interactions with residues in other regions of the upper 50 kDa domain.

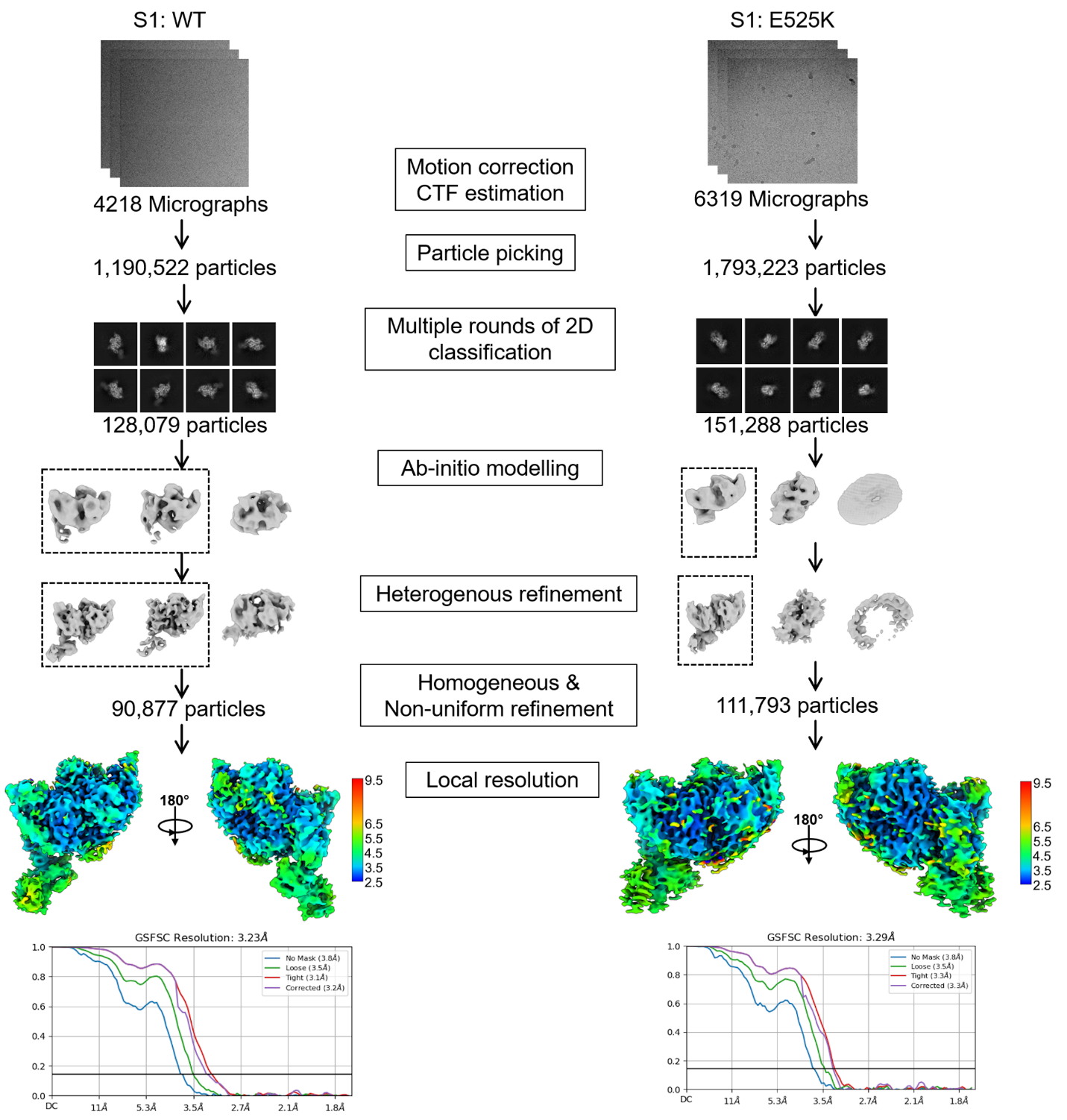

**Fig. S7. Cryo-EM data processing workflow of WT-S1 and E525K-S1.** Schematic overview of the image processing workflow for both WT and E525K S1 structures, including motion correction, CTF estimation, blob and template-based particle picking, multiple rounds of 2D classification to separate junk particles, ab initio reconstruction and heterogeneous refinement for particle curation, homogeneous and non-uniform refinement leads to final reconstruction, colored by local resolution.

**Table S1. Cryo-EM data collection, refinement and validation statistics**

|  | **E525K IHM** | **WT S1** | **E525K S1** |
| --- | --- | --- | --- |
| EMDB ID | EMD-78337 | EMD-78401 | EMD-78403 |
| PDB ID | 37OA | 37PY | 37QC |
| **Data collection and processing** | | |  |
| Magnification | 105,000× | 105,000× | 105,000× |
| Voltage (kV) | 300 | 300 | 300 |
| Pixel size (Å) | 0.83 | 0.83 | 0.83 |
| Defocus range (µm) | 1.0 to 2.0 | 1.0 to 2.0 | 1.0 to 2.0 |
| Total electron dose (e-/Å²) | ~50.0 | ~50.0 | ~50.0 |
| Symmetry | C1 | C1 | C1 |
| No. of movies | 8544 | 4218 | 6319 |
| No. of particles (Final) | 152085 | 90877 | 111793 |
| Map resolution (FSC = 0.143) | 3.0 Å | 3.2 Å | 3.3 Å |
| **Model Refinement and validation** | | |  |
| Initial Model Used (PDB) | 9YOP | 5N69 | 5N69 |
| **Model Composition** | |  |  |
| Non-hydrogen atoms | 19628 | 7363 | 7362 |
| Protein residues | 2435 | 914 | 914 |
| Total ligand molecules | 6 (ADP: 2, PO4: 2, MG: 2) | 3 (ADP: 1, PO4: 1, MG: 1) | 2 (ADP: 1, PO4: 1) |
| **RMSZ deviations** | |  |  |
| Bond length (Å) | 0.17 | 0.17 | 0.29 |
| Bond angle (°) | 0.37 | 0.37 | 0.45 |
| Validation |  |  |  |
| MolProbity Score | 1.87 | 1.78 | 1.93 |
| Clashscore | 11.8 | 10.5 | 12.2 |
| Rotamer Outliers (%) | 0.2 | 0.0 | 0.13 |
| **Ramachandran Plot** | |  |  |
| Favored (%) | 96 | 96.5 | 95.3 |
| Allowed (%) | 4 | 3.5 | 4.7 |
| Disallowed (%) | 0 | 0 | 0 |
| Map-Model CC (overall) | 0.71 | 0.74 | 0.68 |

**Movie S1. 3D variability analysis of the WT IHM along principal component 1 (PC1).**

A series of cryo-EM volumes corresponding to frames 1–20 along PC1 is shown as transparent gray density. The WT docked IHM structure (PDB ID: 9YOP), shown in blue, is fitted into frame 1 to provide structural reference for the conformational changes. The movie illustrates flexibility of the free head (FH) together with the substantial transition of the S2 region from a bent, docked conformation to a straighter, undocked conformation.

**Movie S2. 3D variability analysis of the E525K IHM along principal component 1 (PC1).**

A series of cryo-EM volumes corresponding to frames 1–20 along PC1 is shown as transparent gray density. The E525K IHM structure (PDB ID: 37OA), shown in pink cartoon representation, is fitted into frame 1 to provide structural reference for the observed conformational changes. In contrast to WT, the S2 region remains predominantly bent and docked throughout the PC1 trajectory, whereas the FH retains conformational flexibility.

**Movie S3. MD simulation trajectory of the WT IHM.**

The post-processed and prealigned WT IHM MD trajectory is shown over the 500 ns simulation. The structure is displayed in sphere representation to visualize conformational changes throughout the trajectory. The upper and lower 50-kDa domains are shown in cyan and light blue, respectively; the converter domain is shown in dark blue; the N-terminal domain is shown in gray; The loop 2 regions of both the FH and BH are shown in red and the S2/tail region is shown in purple. The movie highlights the conformational dynamics of the S2 region and Loop 2 regions sampled during the WT simulation.

**Movie S4. MD simulation trajectory of the E525K IHM.**

The post-processed and prealigned E525K IHM molecular dynamics trajectory is shown over the 500 ns simulation. The structure is displayed in sphere representation to visualize conformational changes throughout the trajectory. The upper and lower 50-kDa domains are shown in cyan and light blue, respectively; the converter domain is shown in dark blue; the N-terminal domain is shown in gray; The loop 2 regions of both the FH and BH are shown in red and the S2/tail region is shown in purple. The movie highlights the conformational dynamics of the S2 region and Loop 2 regions sampled during the E525K simulation. This allows comparison of S2 conformational variability observed between WT and E525K trajectories.

**Movie S5. Morph between WT and E525K S1 structures reveals localized mutation-induced conformational changes.**

A structural morph between the WT and E525K S1 cryo-EM structures is shown to visualize the conformational changes associated with the E525K mutation. The E525K S1 structure is colored by per-residue RMSD relative to WT S1, from green (low deviation) to brown (high deviation), highlighting localized structural differences in the activation-loop and Loop 3 region, N-terminal SH3-like domain, and ELC. The morph is played twice in an oscillating manner between the WT and E525K conformations. The view is then rotated by 180°, and the oscillating morph is repeated twice to visualize the conformational changes from the opposite orientation.
